# Adaptive host responses to infection can resemble parasitic manipulation

**DOI:** 10.1101/2023.03.16.532392

**Authors:** Camilla Håkonsrud Jensen, Jacqueline Weidner, Jarl Giske, Christian Jørgensen, Sigrunn Eliassen, Adèle Mennerat

**Affiliations:** Department of Biological Sciences, University of Bergen, Bergen, Norway

**Keywords:** host-parasite coevolution, parasite manipulation, host compensation, hormone strategy, gigantism

## Abstract

Using a dynamic optimisation model for juvenile fish in stochastic food environments, we investigate optimal hormonal regulation, energy allocation and foraging behaviour of a growing host infected by a parasite that only incurs an energetic cost. We find it optimal for the infected host to have higher levels of orexin, growth- and thyroid hormones, resulting in higher activity levels, increased foraging, and faster growth. This growth strategy thus displays several of the fingerprints often associated with parasite manipulation: higher levels of metabolic hormones, faster growth, higher allocation to reserves (i.e. parasite-induced gigantism), higher risk taking and eventually higher predation rate. However, there is no route for manipulation in our model, so these changes reflect adaptive host compensatory responses. Interestingly, several of these changes also increase the fitness of the parasite. Our results call for caution when interpreting observations of gigantism or risky host behaviours as parasite manipulation without further testing.

## Introduction

Hosts and parasites interact antagonistically with each other and many of their traits result from a co-evolutionary arms race (Hudson *et al*. 2006; Brunner *et al*. 2017). In hosts, traits for avoidance of, and resistance against, parasites (see **Table 1** for glossary) are under selection, as evidenced by the wide repertoire of adaptive pre- and post-infection defences. These include reducing infection risk by e.g., avoiding certain areas and types of foods (Hutchings *et al*. 2001), disgust or fear of parasites (Oaten *et al*. 2009; Prokop *et al*. 2010), or prophylactic offspring care (Mennerat *et al*. 2009). Other behaviours occur post-infection, like grooming, behavioural fever, and self-medication (Lefèvre *et al*. 2009; de Roode *et al*. 2013). Hosts can also partly compensate for the detrimental effects of infection via increased foraging effort involving greater risk taking (Milinski 1990; Klein 2003; see also Hite *et al*. 2020). In addition to behavioural defences, organisms have an immune system that protects against and fights infections. Immune defences are costly and often traded-off against other necessary functions such as growth and reproduction (Poulin *et al*. 1994; Sheldon & Verhulst 1996). Hosts may also respond to parasitism by shifting their life histories in adaptive ways e.g., by reproducing earlier in the presence of parasites that strongly compromise future reproduction (Minchella & Loverde 1981; Ebert *et al*. 2004; Gabagambi *et al*. 2020). Finally, if neither resistance nor tolerance of the parasite is possible, host suicide may be adaptive if it increases inclusive fitness (Poulin 1992; Humphreys & Ruxton 2019); infected eusocial insects have for example been observed to move away from their relatives to die in solitude (Heinze & Walter 2010).

**Table 1:** Glossary

|  |  |
| --- | --- |
| **Changes following infection** | Changes in host phenotype (behaviour, physiology, morphology) following a parasitic infection. |
| **Manipulation** | Phenotypic changes in the host induced by parasitic infection that are adaptive for the parasite, but maladaptive for the host. |
| **Compensation** | Adaptive phenotypic changes in the host that compensate for some of the detrimental fitness effects of infection. |
| **(Host) Resistance** | Avoiding or clearing infection. |
| **(Host) Tolerance** | The ability of the infected host to limit the fitness impact of infection. |
| **(Parasite) Exploitation level** | The proportion of the host's energy drained by the parasite, relative to the host's standard metabolic rate (see **Eq. 1**). |
| **Virulence** | The reduction in host fitness that is due to parasitic infection. |

Certain parasites, referred to as manipulative parasites, induce changes in host phenotype that increases their own fitness while being counter-adaptive for the host (Holmes & Bethel 1972; Poulin 1995; Thomas et al. 2005). Host manipulation has been the focus of hundreds of studies and is now recognised as a widespread adaptive strategy for parasites (Poulin & Maure 2015) and one of the best examples of extended phenotype (Dawkins 1982). The changes in host phenotype following infection range from altered host behaviour or morphology resulting in increased predation rates (e.g. *Schistocephalus solidus* infecting copepodites; Hafer & Milinski 2016; changes in eye stalk colouration and shape of snails infected with *Leucochloridium* spp.; Wesołowska & Wesołowski 2014), to gigantism with increased host growth and/or reserves (e.g. *Daphnia magna* infected by *Pasteuria ramosa;* Ebert *et al*. 2004). These modifications can also be accompanied by physiological changes in hormone levels or in the central nervous system of the host (Klein 2003; Escobedo *et al*. 2005).

When host physiology and behaviour change following infection, however, it can sometimes be difficult to assess whether the change is adaptive for the parasite, the host, or is a “by-product” of the infection. The issue fostered decades of research aimed at testing the adaptive consequences of host manipulation for hosts and for parasites (Poulin 2021). Caution is warranted, as appearances can be misleading and only experimental work can allow to disentangle cause from consequence (Poulin & Maure 2015). Besides, most studies of host manipulation have focused on its adaptive value, whereas the underlying proximate mechanisms have largely been overlooked. Identifying the manipulation factors of parasites has been repeatedly called for (Herbison *et al*. 2018; Poulin & Maure 2015); hormones, neurotransmitters, or symbionts are among the proposed candidates (Herbison 2017). For example, infection by the parasitic acanthocephalan *Polymorphus paradoxus* in the gammarid *Gammarus lacustris* leads to increased serotonin levels and associated changes in host phototaxis (Maynard *et al*. 1996; Perrot-Minnot *et al*. 2014). But in most other cases of suspected or established host manipulation there is still a need to establish which pre-existing pathways, within the host, parasites might be exploiting (Lefèvre *et al*. 2009; Helluy & Thomas 2010; Helluy 2013).

In this study, we incorporate current knowledge of the physiological regulation of feeding and juvenile growth of fish in a model, to test (1) whether some of the host phenotypic changes often attributed to parasite manipulation (e.g., higher growth rates, higher risk taking) can arise as adaptive plasticity in the host, as a compensatory response to the energetic costs of parasitism, (2) how optimal host responses to these costs vary according to environmental quality, and (3) whether these changes in the host could also benefit parasites. Using optimisation modelling we start by testing whether the energetic costs of parasitism alone can lead to hormone-mediated increases in host growth, body condition, and exposure to predation. To do so we compare the optimal responses of fish hosts experiencing differing levels of parasite exploitation. By simulating three levels of food availability, we then test how the optimal host responses to parasite exploitation differ across environments. Finally, we explore how parasite exploitation level relates to fitness, either for a parasite still developing in its host or for a trophically-transmitted parasite ready to leave its intermediate host.

## Material and methods

We use an optimisation model of hormonal regulation of growth in fish (Jensen *et al*. 2020a, b; Weidner *et al*. 2020) to study how host growth and behaviour respond to the energetic costs of parasite infection. The model captures the flow of energy through the fish, from foraging, metabolic activities, and digestion to growth, with the endocrine system regulating host energetics and mediating trade-offs with survival. The fish in our model should be seen as juvenile, as for the sake of simplicity we do not consider reproduction or reproductive investment. Here we give a brief explanation of the main features of our model and refer to Weidner *et al*. (2020), Jensen *et al*. (2020a) for further details, including a list of parameters and variables.

One main assumption in the model is that survival (to predation) and physiology are linked via respiration. This approach is built on Priede (1985) as well as empirical studies of the trade-offs between energy acquisition rates and swimming performance in growing Atlantic silversides (*Menidia menidia*, Billerbeck *et al*. 2001; Lankford *et al*. 2001). In the model, we compare the total oxygen use from all aerobic metabolic processes with the maximum oxygen uptake, following Holt & Jørgensen (2014). The more oxygen the fish use relative to maximum oxygen uptake, the less is available for escape, and the more vulnerable the fish will be to predation.

Environments tend to vary gradually, which is often reflected in the fact that current food availability is correlated with that in the near past and future. We incorporate these aspects in our model by adding temporal autocorrelation to food availability. The fish respond to these fluctuations by adjusting their feeding behaviour, growth rate, and metabolism. When the conditions permit it, the fish may build energy reserves that they can draw from in times of scarcity (Jensen *et al*. 2020a).

In the model, we simplify the complex hormonal regulation of feeding and growth to three main functions: The Growth Hormone Function (GHF), the Orexin Function (OXF), and the Thyroid Hormone Function (THF). GHF affects growth rate, OXF appetite, while THF regulates both standard metabolic rate (SMR) and maximum oxygen uptake. For each time step, the model uses stochastic dynamic programming (Houston & McNamara 1999; Clark & Mangel 2000) to maximize host survival until adulthood. It does so by finding the optimal combination of GHF, OXF, and THF for all combinations of two internal and one external state of the fish: stored reserves [J], body length [cm], and food availability [dimensionless].

Hormone levels affect host survival in the following ways (Weidner *et al*. 2020): First, predation risk for fish generally decreases with size, hence more GHF triggering faster growth reduces mortality risk in the long run. Second, fish with higher OXF levels are more actively foraging and thus more exposed to predators. Finally, THF affects mortality in opposite ways by: (1) increasing maximum oxygen uptake, which makes it easier to escape predators, and (2) by increasing metabolic rate, which requires more oxygen and energy, and thus higher foraging activity and risk exposure.

Note that our approach differs from Dynamic Energy Budget (DEB) models in the sense that it explores adaptive changes in growth rates under varying circumstances. For a longer discussion of our approach compared to DEB models, please see Weidner et al. (2020).

### Parasite exploitation of host

In our model we make no assumptions about the life history of the parasite, or whether it is a micro- or macroparasite. Within-host competition is also not explicitly modelled as we make no assumption regarding the number or diversity of parasites infecting the host. For ease of reading, we will here use parasite in the singular form.

The only characteristic of the model parasite is that it takes energy from the host at a certain rate (described below). There is no explicit effect of parasitism on host life history, behaviour, or survival, except that the increased energetic demands due to infection may have knock-on consequences for host mortality, physiology, or behaviour.

The rate at which energy is diverted by the parasite [J min^−1^] is set to be proportional to the metabolic rate of the host:

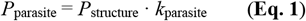

where the coefficient *k*_parasite_ [dimensionless] is the exploitation level of the parasite and *P*_structure_ [J min^−1^] is the structural metabolic rate of the fish. Following Weidner *et al*. (2020) this structural metabolic rate is the product of body mass by an oxygen consumption rate [J min^−1^ g^−1^] under an intermediate level of THF (*τ*_max_/2 [ng ml^−1^] where *τ*_max_ is the maximum THF level [ng ml^−1^]). One of the aims of this study is to compare host responses for different exploitation levels. For the sake of simplicity here these exploitation levels *k*_parasite_ are kept constant throughout each separate simulation.

### Host response to parasites

The model fish has no means of getting rid of the parasite; its only option is to adjust the hormonal regulation of growth and behaviour, ultimately affecting juvenile survival.

Fish may cover the energetic cost of being parasitised by increasing food intake *I* [J min^−1^] or draining energy from reserves *R* [J]. The host’s reserves at the next time step (*t*+1) depend on foraging behaviour and energy allocation in the current time step:

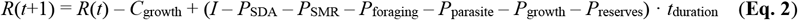

Where *C*_growth_ is the energy incorporated into new structural tissue [J], *I* is intake, *P*_SDA_ is the energetic cost of digesting food [J min^−1^], *P*_SMR_ is the standard metabolic rate under influence of THF [J min^−1^], *P*_foraging_ is the foraging cost [J min^−1^], and *P*_growth_ and *P*_reserves_ are the energetic conversion costs from intake to growth and from reserves to growth [J min^−1^], respectively.

Bioenergetic rates are multiplied by the duration of a time step, *t*_duration_ [min]. Further details can be found in Weidner et al. (2020) were we explore the energetic costs of growth, including conversion costs, in great detail. The only difference between the model presented here and the one used in Weidner *et al*. (2020) and Jensen *et al*. (2020a,b) is the addition of the term *P*_parasite_ representing the rate at which energy is diverted from the host by the parasite **(Eq.2)**.

### Starvation

In addition to mortality due to predation, the model incorporates a negative effect of starvation on host survival. Here host survival *S* [week^−1^] follows a negative exponential that depends on total mortality *M* [year^−1^], as well as on relative energy reserves (*R/R*_max_) and a coefficient of starvation *k*_starvation_ [dimensionless]. If *R* drops below *k*_*s*tarvation_ · *R*_max_ fish survival rapidly declines with relative energy reserves (*R*/*R*_max_):

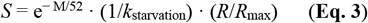

### Experimental simulations

To investigate whether the nature or direction of optimal host responses to parasitism depend on habitat quality, we simulated three groups of individual fish experiencing three different levels of food availability: (1) poor food availability resembling a poor natural environment, (2) intermediate food availability, and (3) rich food availability, where conditions arguably reflect *ad libitum* feeding e.g. in the laboratory. Prior to experimental simulation all individual fish were first optimised to the same wide environmental range of food availabilities spanning all three levels described above.

## Results

The optimal response in fish hosts infected with a parasite diverting energy was to shift hormone levels, which resulted in changes spanning from altered growth rates to modified foraging behaviour and thus exposure to predation.

### Physiological and behavioural changes in the fish host

Fish harbouring parasites with a higher exploitation level experienced higher energetic costs and compensated with increased foraging intensity (**Fig. 1b**). This was a result of elevated appetite, caused by up-regulation of the Orexin Function (OXF) (**Fig. 1e**). Higher parasite exploitation level also increased optimal levels of the Thyroid Hormone Function (THF) (**Fig. 1f**), which in turn led to higher metabolism and increased maximum oxygen uptake.

**Figure 1:**
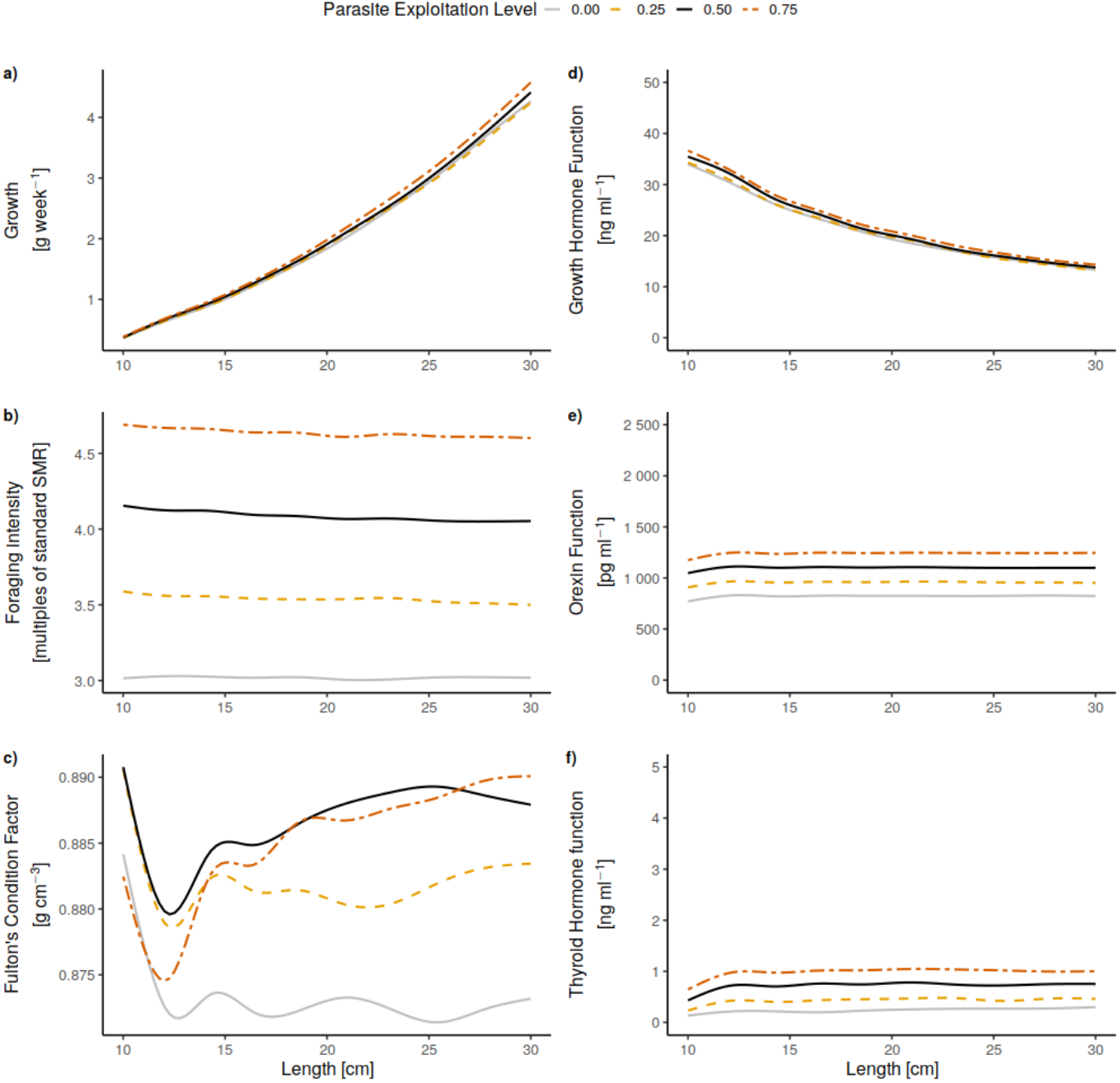
(a) Mean host growth, (b) foraging intensity and (c) Fulton’s condition factor [100 · (*total weight*/*length*^3^)] for different parasite exploitation levels. These emerge from optimising (d) Growth Hormone Function (GHF), (e) Orexin Function (OXF) and (f) Thyroid Hormone Function (THF) levels in our model for each of the four exploitation levels (see Methods for details). Lines are smoothed using a generalised additive model for ease of reading.

Higher foraging intensity and metabolism are expected given the additional energy demand from hosting a parasite. More surprisingly, Growth Hormone Function (GHF) levels and consequently host growth increased with parasite exploitation level (**Fig. 1a & d**, but only in relatively rich environments, see below). Infected hosts also stored more energy in their reserves: At the beginning of the juvenile growth period, the mean Fulton’s condition factor [100 · (*total weight*/*length*^3^)] was higher for hosts infected by parasites with higher exploitation levels, and condition factors increased and stabilised as the fish grew (**Fig. 1c**). Higher condition, foraging activity, metabolism, and growth, however, come at the cost of an increased predation risk (**Fig. 3a**).

### Optimal host strategies under different levels of food availability

In the group that experienced high food availability resembling laboratory conditions (right column of **Fig. 2**), our model predicts faster growth with high-cost parasites. The higher the parasite exploitation level, the faster the host growth, and the higher the mortality risk. These patterns were also found under intermediate food availability (middle column of **Fig. 2**) although the difference among exploitation levels was smaller. In the scenario with poor food availability (left column of **Fig. 2**) the situation was reversed, with heavily parasitised hosts growing more slowly, while taking higher risks when foraging and thus having little chance of surviving.

**Figure 2:**
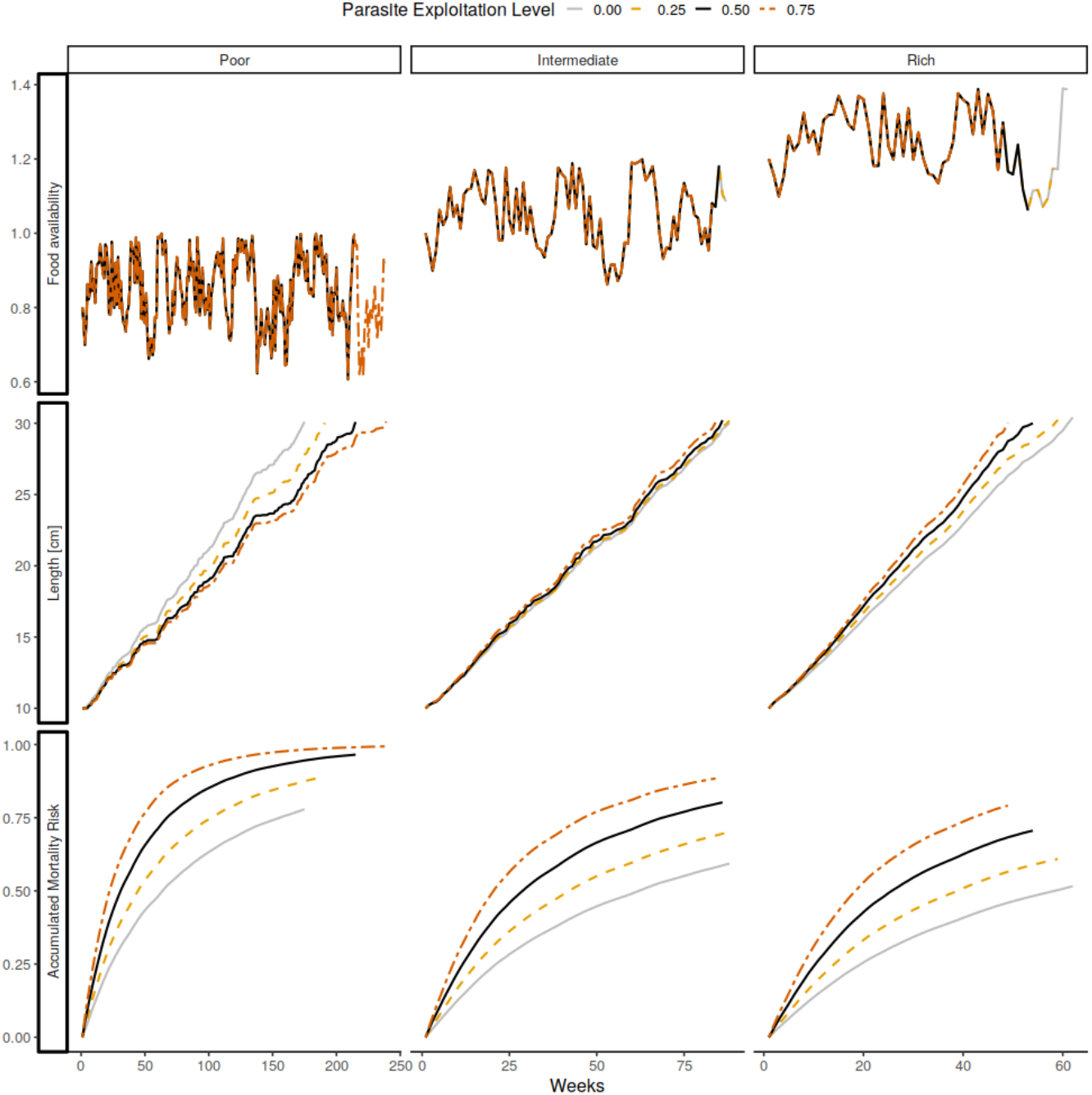
Under conditions of low food availability in the environment (top row), the optimal growth strategy for hosts experiencing high levels of parasite exploitation is to forage more intensely and therefore grow faster (middle row), while the opposite is true in rich environments; mortality is generally higher in the relatively poor environment due to higher foraging (risk-taking), and increases with parasite exploitation level (bottom row).

**Figure 3:**
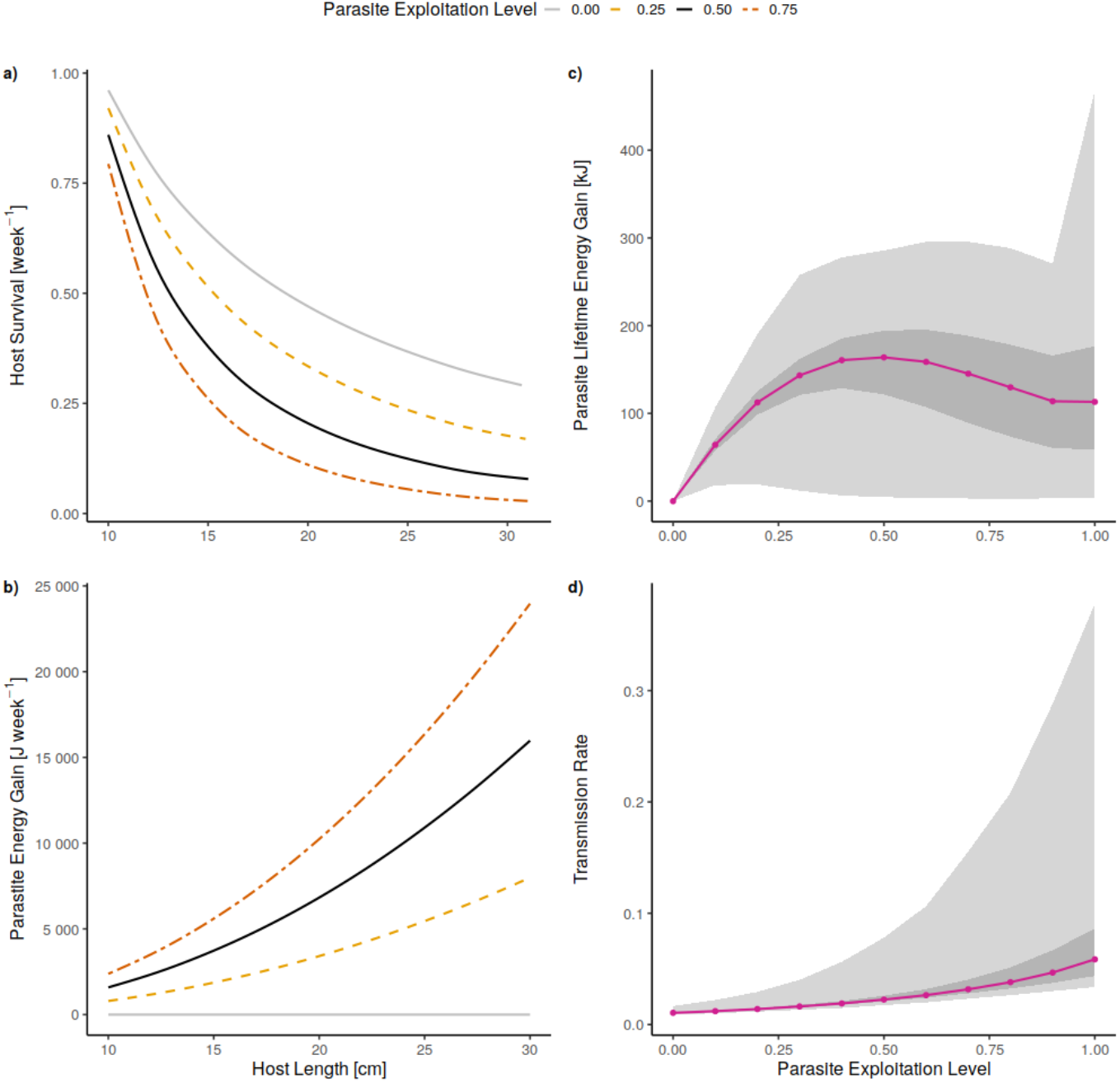
Effects of host responses on proxies of parasite fitness for different exploitation levels. (a) Mean host survival [week^−1^], with predation during foraging being the main cause of mortality in our model; (b) rate of energy gain for the parasite during host growth; (c) Parasite lifetime energy gain (*parasite energy gain* [J week^−1^] · *host survival* [week^−1^]), used here to approximate fitness for a parasite that needs its host to survive. (d) Expected transmission rate (– *log*(*host survival* [week^−1^]) / *host growth period* [weeks]), used here to approximate fitness in those cases where the fish is an intermediate host and the parasite ready to be trophically transmitted to the next host. Violet circles represent median values, dark grey area represent the values from 0.25 to 0.75 quantile, while light grey areas represent the values from 0 to 1 quantile. Lines for a) and b) are smoothed using a generalised additive model for ease of reading.

### Parasites fitness for different exploitation levels, in intermediate or final hosts

Parasite strategies are not optimised in our model, but we explore selection on exploitation levels for parasites at different life stages.

A developing parasite would benefit from not killing its host until it is ready to leave it (in the case of an intermediate host) or have successfully reproduced (in the case of a final host). For such a parasite, lifetime energy gain [kJ] in the host can be used as a fitness proxy. According to our model this proxy for fitness is maximised at an intermediate exploitation level (**Fig. 3c**). In contrast, a trophically-transmitted parasite that is ready to leave its intermediate host would not benefit from letting the host survive, but rather from increasing the probability that the host will be eaten by the next host in its life cycle. Here a more suitable fitness proxy is transmission rate (here defined as – *log*(*host survival* [week^−1^]) / *host growth period* [weeks]), and our model indicates that it increases with exploitation level (**Fig. 3d**).

## Discussion

Here by optimising host responses to parasitism at the hormonal level we find that the optimal response for juvenile parasitised hosts is to increase their feeding- and growth-related hormone levels. The resulting higher foraging intensity, growth, metabolism, and body condition come at the cost of increased predation risk. Furthermore, our model shows that gigantism or increased risk-taking do not only reflect optimal responses in and for the host, but that several of these changes may also benefit the parasite.

Our results align with several former studies showing changes in metabolic rates and performance in infected hosts (Robar *et al*. 2011; Careau *et al*. 2012; Binning *et al*. 2013, 2017; McElroy & de Buron 2014). Increased reserves coupled with growth enhancement may result in gigantism, where hosts increase in size following a parasitic infection. Gigantism has been reported in many taxa, e.g. *Daphnia* (Ebert *et al*. 2004), snails (Ballabeni 1995) and fish (Arnott *et al*. 2000) and is often associated with host castration. According to the temporal storage hypothesis (Ebert *et al*. 2004) host castration benefits the parasite because it keeps the host growing, thereby accumulating reserves that can later be diverted into parasite reproduction. Even though gigantism is often associated with host castration, there are notable exceptions; three-spined sticklebacks (*Gasterosteus aculeatus*) infected by the cestode *Schistocephalus solidus* display increased growth but no reduction in gonadal investment. They are also, like our model fish, heavier than uninfected fish, and show up to 17% increase in the weight of liver reserves (Arnott *et al*. 2000). One explanation may be that enhanced growth is a bet-hedging strategy that helps hosts cope with the risk of starvation. In addition, our results give a hint as to why gigantism is rarely observed in the wild (Fernandez & Esch 1991; Taskinen 1998; Barber *et al*. 2000), as our model only predicts increased growth of infected individuals when food availability is high.

The model described here optimises hormone levels from the perspective of the host only, and not the parasite. Our proxies for parasite fitness (lifetime energy gain or transmission rate), however, indicate that the host responses may also be adaptive for the parasite. The way in which selection favours parasite strategies that best balance extracting energy from the host while keeping it alive (also referred to as the “virulence-transmission trade-off”), has been well-studied in the past decades (e.g. Bull 1994; Jensen *et al*. 2006; Alizon *et al*. 2009; Mennerat *et al*. 2012). Our model also suggests that an intermediate exploitation level is best at solving this trade-off, for parasites with a direct life cycle or for trophically-transmitted parasites in pre-infective stages (**Fig. 3c**). For trophically-transmitted parasites fitness is maximised by exploiting the host as much as possible, inducing risky foraging behaviour, and hence increasing the chances of transmission to the next host (**Fig. 3d**). The fact that host manipulation only occurs at the infective stage is well-described elsewhere; repeatedly measuring hosts and comparing their responses at the pre-*versus* post-infective stage, is commonly used as a way to test whether altered host responses result from manipulation or are mere byproducts (e.g. Poulin 1994; Hafer & Milinski 2015; Gabagambi *et al*. 2019). The novelty here is that our model provides a mechanistic link for how switching from intermediate to high exploitation level as the parasite reaches infective stage may result in corresponding alterations in host behaviour, switching to higher foraging rates involving higher risk-taking and resulting in higher predation rate.

Finally, not all behavioural or physiological changes following infection are explained by host compensatory mechanisms alone. Uncontroversial manipulation of hosts by parasites does exist; insects protecting the pupae of their parasitoids (Libersat *et al*. 2018 and references therein) or “zombie ants” spreading spores of parasitic fungi (Hughes *et al*. 2011) are host manipulation, beyond doubt. Our results show nonetheless that simple physiological mechanisms should be considered as pre-existing paths towards manipulation, and that parasites would be selected for their ability to exploit compensatory responses in hosts whenever those benefit them (Lefèvre *et al*. 2008). Together with earlier studies we argue that the “energy drain hypothesis” and the “parasite manipulation hypothesis” need not be mutually exclusive, and that some unresolved cases might be better understood by adopting a more holistic approach (e.g. Thomas *et al*. 2005; Hafer & Milinski 2016). Behavioural changes following infection, even some of those that in some systems primarily benefit parasites, may in others be adaptive for infected hosts too.

## Acknowledgements

We have been supported by Research Council of Norway (grant number 239834) and the University of Bergen. We thank Knut Helge Jensen, Marc Mangel and Manfred Milinski for discussions and two anonymous referees for feedback on an earlier version. This research contributes to the Centre for Digital Life Norway.

